# Gene model for the ortholog of *DENR* in *Drosophila pseudoobscura*

**DOI:** 10.64898/2026.08.11.744233

**Authors:** Megan E. Lawson, Kylee Sanow, Mihai Fratian, Madelyn Matura, Riley Scanlon, Madeline Richard, Monica Nakhla, Chinmay P. Rele, Jeffrey S. Thompson, Geoffrey D. Findlay, Kellie S. O’Rourke

## Abstract

Gene model for the ortholog of *Density regulated protein* (*DENR*) in the Apr. 2013 (BCM-HGSC Dpse_3.0/DpseGB3) Genome Assembly (GenBank Accession: GCA_000001765.2) of *Drosophila pseudoobscura*. This ortholog was characterized as part of a developing dataset to study the evolution of the Insulin/insulin-like growth factor signaling pathway (IIS) across the genus *Drosophila* using the Genomics Education Partnership gene annotation protocol for Course-based Undergraduate Research Experiences.

## Introduction

“Computational gene predictions in non-model organisms often can be improved by careful manual annotation and curation, allowing for more accurate analyses of gene and genome evolution (Mudge and Harrow 2016; Tello-Ruiz et al., 2019). The Genomics Education Partnership (thegep.org) uses web-based tools to allow undergraduates to participate in course-based research by generating manual annotations of genes in non-model species (Rele et al., 2023). These models of orthologous genes across species, such as the one presented here, then provide a reliable basis for further evolutionary genomic analyses when made available to the scientific community. The particular gene ortholog described here, *Density regulated protein* (*DENR*) in *D. pseudoobscura*, was characterized as part of a developing dataset to study the evolution of the Insulin/insulin-like growth factor signaling pathway (IIS) across the genus Drosophila.” (Myers et al., 2024).

“The IIS pathway is a highly conserved signaling pathway in animals and is central to mediating organismal responses to nutrients (Hietakangas and Cohen 2009; Grewal 2009)” (Myers et al., 2024). “*DENR* was first discovered in a human teratocarcinoma cell line because its concentration in cells increased with cell density (Deyo et al., 1998). Subsequent bioinformatic and biochemical analyses showed that the protein is conserved across eukaryotes and functions in non-canonical translation initiation (Fleischer et al., 2006; Skabkin et al., 2010). *D. melanogaster* flies homozygous for a null, knockout allele of the gene encoding *DENR* (FBgn0030802), die as pharate adults, showing a larval-like epidermis and reduced proliferation of histoblast cells (Schleich et al., 2014). Subsequent experiments using both RNAi in S2 cells and the knockout allele in larvae showed that DENR is required, along with its interacting partner MCT-1, for the proper expression regulation of a subset of transcripts required for cell cycle progression and growth. In particular, the loss of *DENR* reduces expression of the insulin receptor and makes larvae less sensitive to insulin signaling (Schleich et al., 2014), thus implicating DENR in the regulation of the insulin signaling pathway.” (Laskowski et al., 2024).

“*D. pseudoobscura* is part of the *pseudoobscura* species subgroup within the *obscura* species group in the subgenus *Sophophora* of the genus *Drosophila* (Sturtevant 1942; Buzzati-Traverso and Scossiroli 1955). It was first described by Frolowa and Astaurow (1929). The *pseudoobscura* species subgroup is endemic to the western hemisphere, where *D. pseudoobscura* is distributed throughout Western North America, Mexico, and Central America (Markow and O’Grady 2005). An additional population of *D. pseudoobscura*, found near Bogota, Colombia, is partially reproductively isolated from the North and Central American populations (Prakash & Merritt 1972). *D. pseudoobscura* is found primarily in chaparral and temperate forests. *D. pseudoobscura* has been studied extensively in the context of ecological and behavioral genetics, speciation, and genome evolution (Powell 1997).” (Lawson et al., 2025).

We propose a gene model for the *D. pseudoobscura* ortholog of the *D. melanogaster Density regulated protein* (*DENR*) gene. The genomic region of the ortholog corresponds to the uncharacterized protein XP_001354336.1 (Locus ID LOC4814209) in the Apr. 2013 (BCM-HGSC Dpse_3.0/DpseGB3) Genome Assembly of *D. pseudoobscura* (GenBank Accession: GCA_000001765.2). This model is based on RNA-Seq data from *D. pseudoobscura* (Chen et al. 2014; SRP006203) and *DENR* in *D. melanogaster* using FlyBase release FB2024_02 (GCA_000001215.4; Gramates et al., 2022; Jenkins et al., 2022; Larkin et al., 2021). The Genomics Education Partnership maintains a mirror of the UCSC Genome Browser (Kent WJ et al., 2002; Gonzalez et al., 2021), which is available at https://gander.wustl.edu.

## Results

### Synteny

The target gene, *DENR*, occurs on chromosome X in *D. melanogaster* and is flanked upstream by *CG4880* and *CG13002* and downstream by *RNA polymerase III subunit I* (*Polr3I*) and *Nitrogen permease regulator-like 2* (*Nprl2*). The *tblastn* search of *D. melanogaster* DENR-PA (query) against the *D. pseudoobscura* (GenBank Accession: GCA_000001765.2) Genome Assembly (database) placed the putative ortholog of *DENR* within scaffold CH379066 (CH379066.3) at locus LOC4814209 (XP_001354336.1), with an e-value of 2e-56 and a percent identity of 61.49%. The putative ortholog is flanked upstream by LOC4814194 (XP_001354337.1) and LOC4814197 (XP_001354338.3), which correspond to *Nprl2* and *CG4872* in *D. melanogaster* (e-values: 0.0 and 1e-162; percent identities: 92.23% and 60.97%, respectively, as determined by *blastp*; Figure 1A, Altschul et al., 1990). The putative ortholog of *DENR* is flanked downstream by LOC4814215 (XP_001354335.2) and LOC4814191 (XP_001354334.2), which correspond to *Polr3I* and *CG4880* in *D. melanogaster* (e-values: 6e-81 and 2e-73; percent identities: 68.39% and 43.38%, respectively, as determined by *blastp*) (**Figure 1A**). The putative ortholog assignment for *DENR* in *D. pseudoobscura* is supported by the following evidence: synteny of the genomic neighborhood is partially conserved, and additionally while the locations of *Nprl2* and *CG4880* are not syntenic, they both remain in the genomic neighborhood of *DENR*. All *blast* results used to determine orthology indicate very high quality matches.

**Figure 1.**
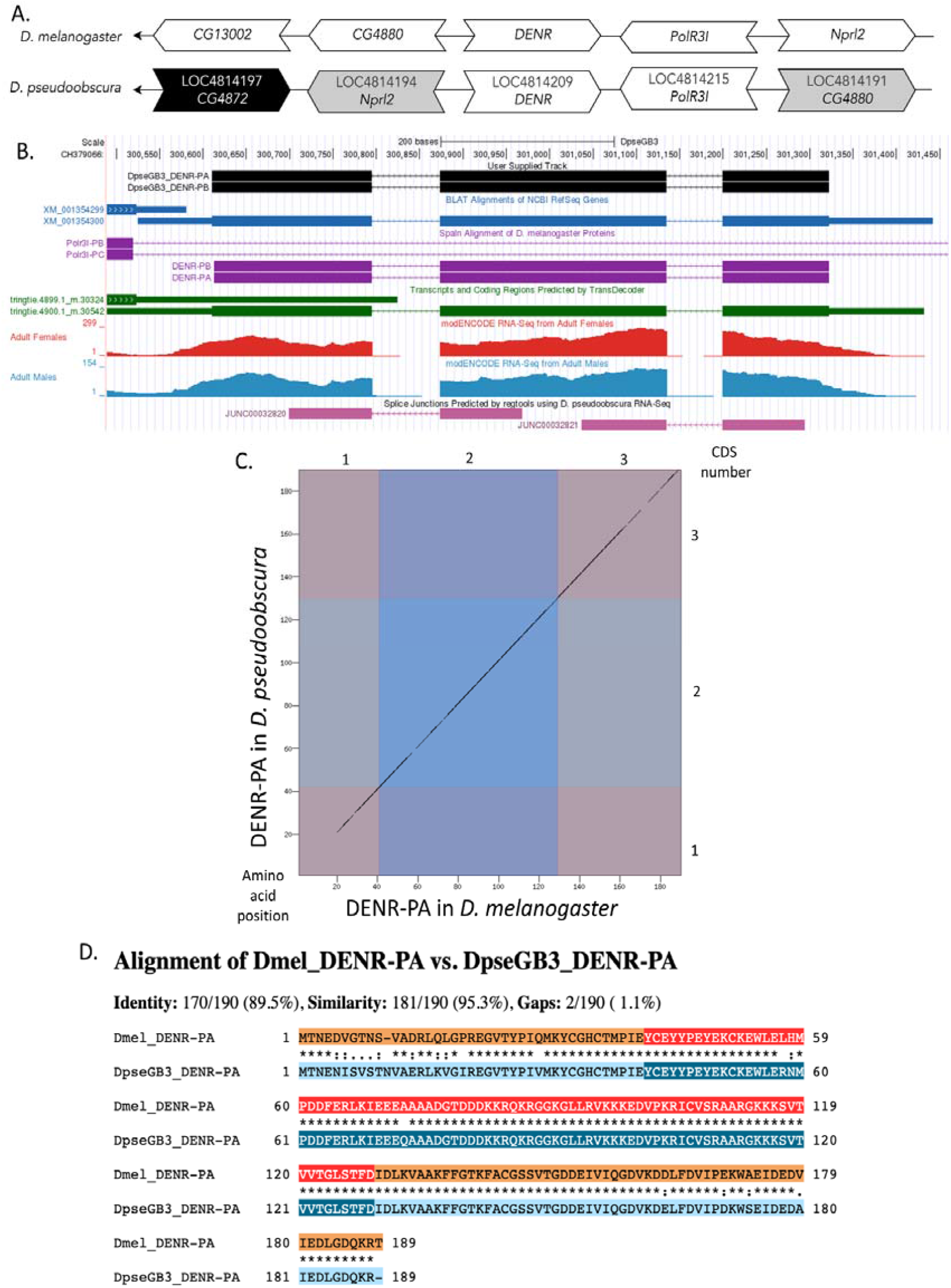
(A) Synteny comparison of the genomic neighborhoods for *DENR* in *Drosophila melanogaster* and *D. pseudoobscura*. Thin underlying arrows indicate the DNA strand within which the target gene–*DENR*–is located in *D. melanogaster* (top) and *D. pseudoobscura* (bottom). Thin arrows pointing to the left indicate that *DENR* is on the negative (-) strand in *D. melanogaster* and *D. pseudoobscura*. The wide gene arrows pointing in the same direction as *DENR* are on the same strand relative to the thin underlying arrows, while wide gene arrows pointing in the opposite direction of *DENR* are on the opposite strand relative to the thin underlying arrows. White gene arrows in *D. pseudoobscura* indicate orthology to the corresponding gene in *D. melanogaster*, black gene arrows indicate non-orthology, and gray gene arrows indicate that a gene is present in both neighborhoods but in different locations relative to the target gene. Gene symbols given in the *D. pseudoobscura* gene arrows indicate the orthologous gene in *D. melanogaster*, while the locus identifiers are specific to *D. pseudoobscura*. **(B) Gene Model in GEP UCSC Track Data Hub** (Raney et al., 2014). The coding-regions of *DENR* in *D. pseudoobscura* are displayed in the User Supplied Track (black); coding CDSs are depicted by thick rectangles and introns by thin lines with arrows indicating the direction of transcription. Subsequent evidence tracks include BLAT Alignments of NCBI RefSeq Genes (dark blue, alignment of Ref-Seq genes for *D. pseudoobscura*), Spaln of D. melanogaster Proteins (purple, alignment of Ref-Seq proteins from *D. melanogaster*), Transcripts and Coding Regions Predicted by TransDecoder (dark green), RNA-Seq from Adult Females and Adult Males (red and light blue, respectively; alignment of Illumina RNA-Seq reads from *D. pseudoobsura*), and Splice Junctions Predicted by regtools using *D. pseudoobscura* RNA-Seq (Chen et al., 2014; SRP006203). Each splice junction is supported by 100-499 reads, which is indicated in pink. **(C) Dot Plot of DENR-PA in *D. melanogaster* (*x*-axis) vs. the orthologous peptide in *D. pseudoobscura* (*y*-axis)**. Amino acid number is indicated along the left and bottom; CDS number is indicated along the top and right, and CDSs are also highlighted with alternating colors. The gap in the dot plot region corresponding to the first CDS of DENR-PA results from decreased sequence identity in this region. **(D) Protein alignment of DENR-PA in *D. melanogaster* (top row) vs. the orthologous peptide in *D. pseudoobscura* (bottom row)**. The alternating colored rectangles represent adjacent CDS. The symbols in the match line denote the level of similarity between the aligned residues. An asterisk (*) indicates that the aligned residues are identical. A colon (:) indicates the aligned residues have highly similar chemical properties—roughly equivalent to scoring > 0.5 in the Gonnet PAM 250 matrix (Gonnet et al., 1992). A period (.) indicates that the aligned residues have weakly similar chemical properties—roughly equivalent to scoring > 0 and ≤ 0.5 in the Gonnet PAM 250 matrix. A space indicates a gap or mismatch when the aligned residues have a complete lack of similarity— roughly equivalent to scoring ≤ 0 in the Gonnet PAM 250 matrix. This protein alignment shows that while there is decreased sequence similarity within the beginning portion of the first CDS of DENR-PA in *D. melanogaster* and *D. pseudoobscura*, there is still high conservation of the peptide sequence and the biochemical properties of this region of the peptide.

### Protein Model

*DENR* in *D. pseudoobscura* has three CDSs within the genome sequence. The single unique protein sequence (DENR-PA and DENR-PB) is translated from two mRNA isoforms that differ in their UTRs (*DENR-RA* and *DENR-RB*; **Figure 1B**). Relative to the ortholog in *D. melanogaster*, the CDS number and protein isoform count are conserved, as *DENR-RA* and *DENR-RB* are also identical with three coding CDSs in *D. melanogaster*. The sequence of DENR-PA in *D. pseudoobscura* has 89.5% identity with the protein-coding isoform DENR-PA in *D. melanogaster*, as determined by *blastp* (**Figure 1C**). Reduced sequence identity was found in the first CDS, but the protein alignment (**Figure 1D**) indicates that this region still has very high biochemical similarity across the two species. Coordinates of this curated gene model (DENR-PA, DENR-PB) are stored by NCBI at GenBank/BankIt (accession BK064553, BK064554, respectively). These data are also archived in the CaltechDATA repository (see “Extended Data” section below).

## Methods

Detailed methods including algorithms, database versions, and citations for the complete annotation process can be found in Rele et al. (2023).

## Acknowledgements

We would like to thank Wilson Leung for developing and maintaining the technological infrastructure that was used to create this gene model and Laura K. Reed for overseeing the project.

## Supplemental Files

1. Zip file containing FASTA, PEP, GFF files for the gene model

## Metadata

Bioinformatics, Genomics, *Drosophila*, Genotype Data, New Finding

## References

Altschul, S. F., Gish, W., Miller, W., Myers, E. W., & Lipman, D. J. (1990). Basic local alignment search tool. Journal of Molecular Biology, 215(3), 403–410. 10.1016/S0022-2836(05)80360-2

Buzzati-Traverso, A. A., & Scossiroli, R. E. (1955). The “Obscura Group” of the Genus Drosophila. In Advances in Genetics (Vol. 7, pp. 47–92). Elsevier. 10.1016/S0065-2660(08)60093-0

Chen, Z.-X., Sturgill, D., Qu, J., Jiang, H., Park, S., Boley, N., Suzuki, A. M., Fletcher, A. R., Plachetzki, D. C., FitzGerald, P. C., Artieri, C. G., Atallah, J., Barmina, O., Brown, J. B., Blankenburg, K. P., Clough, E., Dasgupta, A., Gubbala, S., Han, Y., … Richards, S. (2014). Comparative validation of the D. melanogaster modENCODE transcriptome annotation. Genome Research, 24(7), 1209–1223. 10.1101/gr.159384.113

Deyo, J. E., Chiao, P. J., & Tainsky, M. A. (1998). Drp, a Novel Protein Expressed at High Cell Density but Not During Growth Arrest. DNA and Cell Biology, 17(5), 437–447. 10.1089/dna.1998.17.437

Fleischer, T. C., Weaver, C. M., McAfee, K. J., Jennings, J. L., & Link, A. J. (2006). Systematic identification and functional screens of uncharacterized proteins associated with eukaryotic ribosomal complexes. Genes & Development, 20(10), 1294–1307. 10.1101/gad.1422006

Frolowa, S. L., & Astaurow, B. L. (1929). Die Chromosomengarnitur als systematisches Merkmal: Eine vergleichende Untersuchung der russischen und amerikanischen Drosophila obscura Fall. Zeitschrift für Zellforschung und Mikroskopische Anatomie, 10(1), 201–213. 10.1007/BF02450642

Gonzalez, J. N., Zweig, A. S., Speir, M. L., Schmelter, D., Rosenbloom, K. R., Raney, B. J., Powell, C. C., Nassar, L. R., Maulding, N. D., Lee, C. M., Lee, B. T., Hinrichs, A. S., Fyfe, A. C., Fernandes, J. D., Diekhans, M., Clawson, H., Casper, J., Benet-Pagès, A., Barber, G. P., … Kent, W. J. (2021). The UCSC Genome Browser database: 2021 update. Nucleic Acids Research, 49(D1), D1046– D1057. 10.1093/nar/gkaa1070

Gramates, L. S., Agapite, J., Attrill, H., Calvi, B. R., Crosby, M. A., Dos Santos, G., Goodman, J. L., Goutte-Gattat, D., Jenkins, V. K., Kaufman, T., Larkin, A., Matthews, B. B., Millburn, G., Strelets, V. B., the FlyBase Consortium, Perrimon, N., Gelbart, S. R., Agapite, J., Broll, K., … Lovato, T. (2022). FlyBase: A guided tour of highlighted features. Genetics, 220(4), iyac035. 10.1093/genetics/iyac035

Grewal, S. S. (2009). Insulin/TOR signaling in growth and homeostasis: A view from the fly world. The International Journal of Biochemistry & Cell Biology, 41(5), 1006–1010. 10.1016/j.biocel.2008.10.010

Hietakangas, V., & Cohen, S. M. (2009). Regulation of Tissue Growth through Nutrient Sensing. Annual Review of Genetics, 43(1), 389–410. 10.1146/annurev-genet-102108-134815

Jenkins VK, Larkin A, Thurmond J. Using FlyBase, a database of Drosophila gnes and genetics. In: Dahmann C, editor. Drosophila: methods and Protocols. New York (NY): Springer; 2022.

Kent, W. J., Sugnet, C. W., Furey, T. S., Roskin, K. M., Pringle, T. H., Zahler, A. M., & Haussler, A. D. (2002). The Human Genome Browser at UCSC. Genome Research, 12(6), 996–1006. 10.1101/gr.229102

Larkin, A., Marygold, S. J., Antonazzo, G., Attrill, H., dos Santos, G., Garapati, P. V., Goodman, J. L., Gramates, L. S., Millburn, G., Strelets, V. B., Tabone, C. J., Thurmond, J., FlyBase Consortium, Perrimon, N., Gelbart, S. R., Agapite, J., Broll, K., Crosby, M., Dos Santos, G., … Lovato, T. (2021). FlyBase: Updates to the Drosophila melanogaster knowledge base. Nucleic Acids Research, 49(D1), D899–D907. 10.1093/nar/gkaa1026

Laskowski, L. F., Burton, I., Stanek, T. J., Findlay, G. D., Tanner, S., Vincent, J. A., Chak, S. T., Ellison, C. E., & Rele, C. P. (2024). Gene model for the ortholog of DENR in Drosophila yakuba. 10.17912/micropub.biology.001017

Lawson M.E., Dufur R, Wright B, Wellik IG, Long LJ, Thompson JS, Rele CP. 2025. Gene model for the ortholog of Roc1a in Drosophila pseudoobscura. Manuscript submitted.

Markow, T. A., & O’Grady, P. M. (2005). Evolutionary Genetics of Reproductive Behavior in Drosophila: Connecting the Dots. Annual Review of Genetics, 39(1), 263–291. 10.1146/annurev.genet.39.073003.112454

Mudge, J. M., & Harrow, J. (2016). The state of play in higher eukaryote gene annotation. Nature Reviews Genetics, 17(12), 758–772. 10.1038/nrg.2016.119

Myers A., Hoffmann A., Natysin M., Arsham A.M, Stamm J., Thompson J.S., Rele C.P. 2024. Gene model for the ortholog Myc in Drosophila ananassae, microPublication Biology, submitted.

Powell, J. R. (1997). Progress and Prospects in Evolutionary Biology: The Drosophila Model. Oxford University Press New York, NY. 10.1093/oso/9780195076912.001.0001

Prakash, S., & Merritt, R. B. (1972). DIRECT EVIDENCE OF GENIC DIFFERENTIATION BETWEEN SEX RATIO AND STANDARD GENE ARRANGEMENTS OF X CHROMOSOME IN DROSOPHILA PSEUDOOBSCURA. Genetics, 72(1), 169–175. 10.1093/genetics/72.1.169

Raney, B. J., Dreszer, T. R., Barber, G. P., Clawson, H., Fujita, P. A., Wang, T., Nguyen, N., Paten, B., Zweig, A. S., Karolchik, D., & Kent, W. J. (2014). Track data hubs enable visualization of user-defined genome-wide annotations on the UCSC Genome Browser. Bioinformatics, 30(7), 1003–1005. 10.1093/bioinformatics/btt637

Rele, C. P., Sandlin, K. M., Leung, W., & Reed, L. K. (2023). Manual annotation of Drosophila genes: A Genomics Education Partnership protocol. F1000Research, 11, 1579. 10.12688/f1000research.126839.3

Schleich, S., Strassburger, K., Janiesch, P. C., Koledachkina, T., Miller, K. K., Haneke, K., Cheng, Y.-S., Küchler, K., Stoecklin, G., Duncan, K. E., & Teleman, A. A. (2014). DENR–MCT-1 promotes translation re-initiation downstream of uORFs to control tissue growth. Nature, 512(7513), 208–212. 10.1038/nature13401

Skabkin, M. A., Skabkina, O. V., Dhote, V., Komar, A. A., Hellen, C. U. T., & Pestova, T. V. (2010). Activities of Ligatin and MCT-1/DENR in eukaryotic translation initiation and ribosomal recycling. Genes & Development, 24(16), 1787–1801. 10.1101/gad.1957510

Sturtevant, A. H. 1942. The classification of the genus Drosophila with the description of nine new species. Univ. Texas Publ. 4213, 5–51

Tello-Ruiz, M. K., Marco, C. F., Hsu, F.-M., Khangura, R. S., Qiao, P., Sapkota, S., Stitzer, M. C., Wasikowski, R., Wu, H., Zhan, J., Chougule, K., Barone, L. C., Ghiban, C., Muna, D., Olson, A. C., Wang, L., Ware, D., & Micklos, D. A. (2019). Double triage to identify poorly annotated genes in maize: The missing link in community curation. PLOS ONE, 14(10), e0224086. 10.1371/journal.pone.0224086

